# Biochemical, structural insights of newly isolated AA16 family of Lytic Polysaccharide Monooxygenase (LPMO) from *Aspergillus fumigatus* and investigation of its synergistic effect using biomass

**DOI:** 10.1101/2020.04.24.059154

**Authors:** Musaddique Hossain, Subba Reddy Dodda, Bishwajit Singh Kapoor, Kaustav Aikat, Sudit S. Mukhopadhyay

**Affiliations:** Department of Biotechnology, National Institute of Technology Durgapur-713209, West Bengal, India

**Keywords:** *A.fumigatus*, Auxiliary activity, Cloning, Kinetics, LPMO, Lignocelluloses, Molecular docking

## Abstract

The efficient conversion of lignocellulosic biomass into fermentable sugar is a bottleneck for the cheap production of bio-ethanol. The recently identified enzyme Lytic Polysaccharide Monooxygenase (LPMO) family has brought new hope because of its boosting capabilities of cellulose hydrolysis. In this report, we have identified and characterized a new class of auxiliary (AA16) oxidative enzyme LPMO from the genome of a locally isolated thermophilic fungus *Aspergillus fumigatus* (NITDGPKA3) and evaluated its boosting capacity of biomass hydrolysis. The *Af*LPMO16 is an intronless gene and encodes the 29kDa protein. While Sequence-wise, it is close to the C1 type of *Aa*AA16 and cellulose-active AA10 family of LPMOs, but the predicted three-dimensional structure shows the resemblance with the AA11 family of LPMO (PDB Id: 4MAH). The gene was expressed under an inducible promoter (AOX1) with C-terminal His tag in the *Pichia pastoris*. The protein was purified using Ni-NTA affinity chromatography, and we studied the enzyme kinetics with 2,6-dimethoxyphenol. We observed polysaccharides depolymerization activity with Carboxymethyl cellulose (CMC) and Phosphoric acid swollen cellulose (PASC). Moreover, the simultaneous use of cellulase cocktail (commercial) and *Af*LPMO16 enhances lignocellulosic biomass hydrolysis by 2-fold, which is highest so far reported in the LPMO family.

**Importance:** The auxiliary enzymes, such as LPMOs, have industrial importance. These enzymes are used in cellulolytic enzyme cocktail due to their synergistic effect along with cellulases. In our study, we have biochemically and functionally characterized the new AA16 family of LPMO from *Aspergillus fumigatus* (NITDGPKA3). The biochemical characterization is the fundamental scientific elucidation of the newly isolated enzyme. The functional characterization, biomass degradation activity of *Af*LPMO16, and cellulase cocktail (commercial) combination enhancing the activity by 2-fold. This enhancement is the highest reported so far, which gives the enzyme *Af*LPMO16 enormous potential for industrial use.

## Introduction

The diminution of fossil fuels and the growing concern of environmental consequences, particularly climate changes, have steered our fast-growing economy for clean and renewable energy production [1]. Among different renewable energy sources, bioethanol is one of the promising alternatives to fossil fuel because of its low CO_2_ emission [2, 3] and its manufacturing reliance on lignocellulosic biomass, which is bio-renewable and abundance on earth. However, the structural complexity and the recalcitrance of this renewable carbon source [4] have hindered its optimal use. The current process of saccharification of lignocellulosic biomass is time-consuming and costly. Therefore, the requirement of cost-effective and fast controlled destruction of lignocellulose has driven the bioethanol industry to explore the accessory enzymes to achieve a better and efficient enzyme cocktail for the commercial production of lignocellulose-derived ethanol.

A breakthrough in such exploration came into existence when a mono-copper redox enzyme, known as Lytic polysaccharide monooxygenase (LPMO), was first reported in 2010 [5-8]. LPMO increases lignocellulosic biomass conversion efficiency[9,10] by catalyzing the hydroxylation of C1 and/or C4 carbon involved in glycosidic bonds that connect glucose unit in cellulose and allow cellulase enzymes to process the destabilized complex polysaccharides [11-15]. Harris et al., in their study, used LPMO from *T reesei* along with classical cellulases and showed that the degradation of polysaccharide substrates was increased by a factor of two when compared with the activity of classical cellulases alone [16]. A CBM33 domain-containing enzyme identified from *Serratia marcescens* with boosting chitinase activity, later classified as LPMO. A study by Nakagawa et al. showed that an AA10 family of LPMO from *Streptomyces griseus* could increase the efficiency of chitinase enzymes by 30- and 20-fold on both α and β forms of chitin, respectively [17]. Along with this work, there are some recent reports of the synergistic effect of LPMOs with glycoside hydrolases on polysaccharide substrates [18-20].

LPMOs are classified as AA9, AA10, AA11, AA13, AA14, and AA15 in the CAZy database (http://www.cazy.org/), based on their amino acid sequence similarity. Recently Filiatrault-Chastel et al. identified the AA16, a new family of LPMO from the secretome of a fungi *Aspergillus aculeatus* (*Aa*AA16). The *Aa*AA16 was initially isolated as X273 protein (unnamed domain) and later identified as C1-oxidizing LPMO active on cellulose [21]. *Aa*AA16, the only AA16 family of LPMO so far, has been identified, and it lacks complete biochemical characterization. The biochemical characterization, structural characterization, and the assessment of biomass conversion efficiency are required to understand better the action of members of this new family on plant biomass and their possible biological roles.

While we were analyzing the cellulose hydrolyzing genes from the genome of *A. fumigatus* (*Aspergillus* genome database), we identified five LPMOs, one belonging to AA16 family because of its X273 domain. Further, we cloned the *Af*LPMO16 gene from the genome of our locally isolated strain of *A. fumigatus* (NITDGPKA3) [22] (GenBank accession No. JQ046374) by designing the primers based on the *A. fumigatus* LPMO sequence (CAF32158.1)(NCBI). The cloned *A. fumigatus* (NITDGPKA3) LPMO (after cloning and sequencing the sequence submitted to GenBank; accession No. MT462230) is expressed in *Pichia pastoris* X33. The heterologous protein (*Af*LPMO16) purified and used for biochemical and functional characterization. The saccharification rate assessment suggests that *Af*LPMO16 has fast and effective glucose releasing ability from lignocellulose and cellulose when used with a commercial cellulase cocktail. Enzyme kinetics using 2,6-dimethoxyphenol as a substrate [23] confirmed the oxidative activity. The lignocellulosic biomass (alkaline pre-treated raw rice straw) conversion efficiency along with cellulases suggests that *Af*LPMO16 could be an essential member of the cellulase cocktail for industrial use.

## Results

### Cloning, expression, and purification of AfLPMO16

*Af*LPMO16 (GenBank accession No. MT462230) is an intronless 870 nucleotide long gene that encodes 290 amino acids. The theoretical molecular mass is 29KDa (including signal peptide). The gene sequence of AA16 from our isolated strain of *A*.*fumigatus* (NITDGPKA3) has shown almost 99.6% homology with the gene sequence of AA16 present in the genome database of *A*.*fumigatus* (CAF32158.1) (NCBI database).

The protein of *Af*LPMO16 (GenBank accession No. MT462230) was produced in *Pichia pastoris* X33 without its C-terminal extension. After the optimization of the expression procedure, we achieved approximately 0.8 mg/ml of purified protein. The SDS-PAGE analysis (Fig 1) confirmed the single band of the purified protein (Fig. 1: lanes 5 and 6). We further confirmed the purified recombinant protein bearing the 6X His-tag by Western blot using an anti-His antibody (Fig. 1: Lane W1 & W2); the purified protein (lane 5 & 6 of SDS-PAGE) used for western blot. The expressed recombinant *Af*LPMO16 band appeared at approximately 32kDa position in SDS-PAGE (Fig. 1), which is slightly higher than the expected size. It is probably due to glycosylation [24], or recombinant protein has *c-myc* epitope and 6x His tag in its c-terminal that can increase the molecular mass by 2.7KDa. For further confirmation of N-glycosylation, we checked the *Af*LPMO16 sequence glycosylation site using NetNGlyc 1.0 server (DTU Bioinformatics, Technical University of Denmark, http://www.cbs.dtu.dk/services/NetNGlyc/) [36]. There were two N-glycosylation sites present above the 0.5 threshold value at 114 & 149 amino acid sequence positions with 0.76 and 0.56 potential values, respectively.

**Figure 1.**
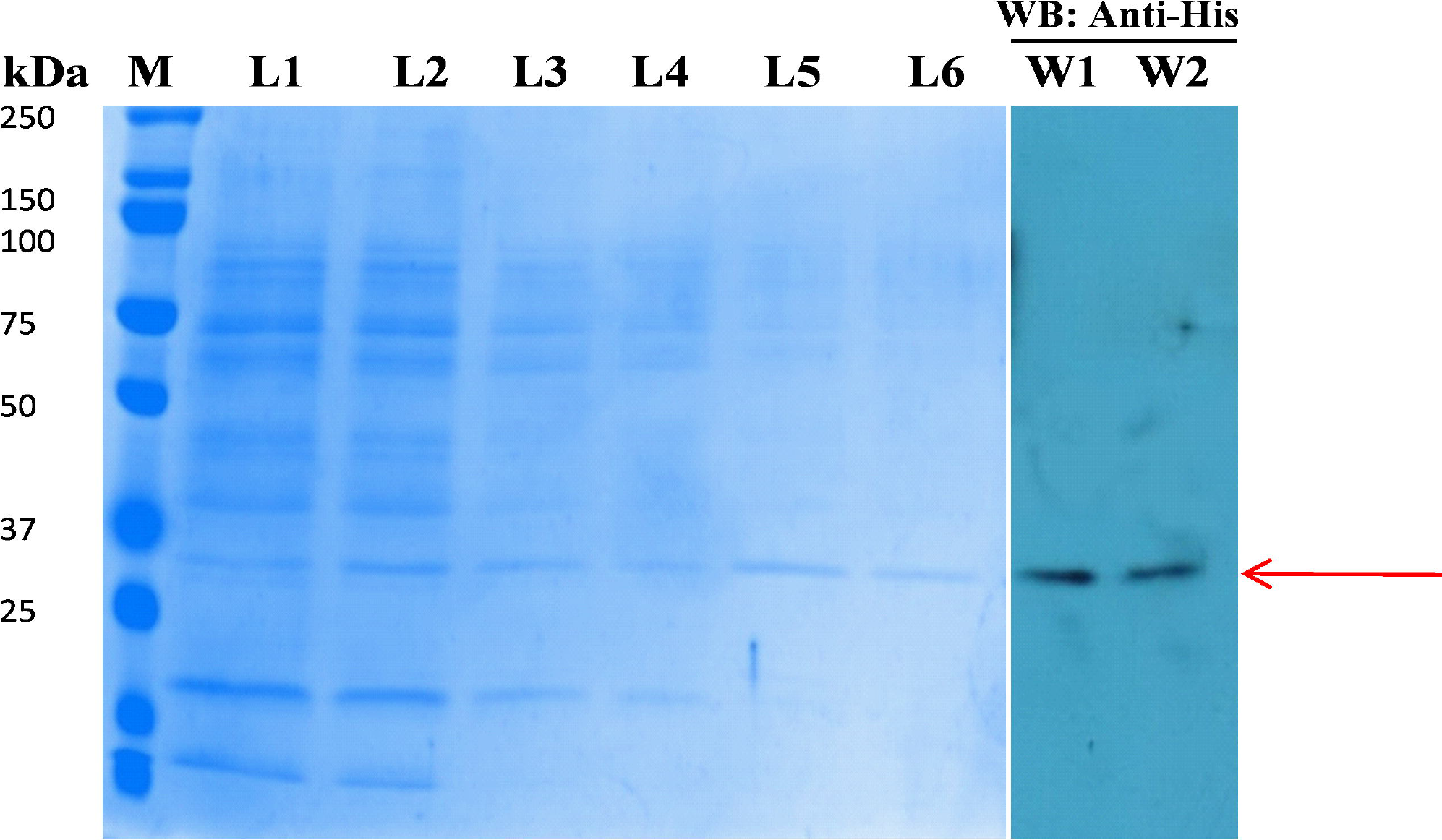
Expression and purification of *Af*LPMO16 (marked with red arrow). SDS PAGE analysis; lane1, flow-through, lane2,3&4 wash, lane 5 & 6. Purified *Af*LPMO16: Western blot analysis using purified protein presented in lane 5 & 6 of SDS page marked as lane W1 and W2

### Enzyme assay and Kinetics

LPMO converts 2,6-dimethoxyphenol (2,6-DMP) into 1-Coerulignone (Fig. 2a) due to its oxidative property, and 1-Coerulignone has an extinction coefficient of 53200. 1-Coerulignone gives absorbance at 469nm wavelength; therefore, we can easily quantify it using a spectrophotometer [21]. The OD at 469nm wavelength steadily increases with time that clearly indicates the steady conversion of 2,6-dimethoxyphenol to 1-Coerulignone (Fig. 2a). It also suggests the sufficient activity of the enzyme *Af*LPMO16. Temperature and pH influence the activity of LPMO. Thus, during the kinetic study, we used optimum temperature 30 and pH 6.0, as described by [21]. *Af*LPMO16 showed proper activity for the chemical substrate 2,6-dimethoxyphenol; there was a steady release of 1-Coerulignone when incubated 2,6-dimethoxyphenol with *Af*LPMO16. The enzyme kinetics was performed with different concentrations of 2,6-dimethoxyphenol. We obtained the Kinetics parameters such as Michaelis Menten constant (K_m_) and maximum velocity (V_max_) from the Line-weaver-Burk plot (Fig. 2b) as 5.4mM, and 0.153 U/mg, respectively. The calculated catalytic activity K_cat_ was 277.67 min^-1^ (Table 1). These kinetics parameters suggest that the oxidative property of *Af*LPMO16.

**Figure 2.**
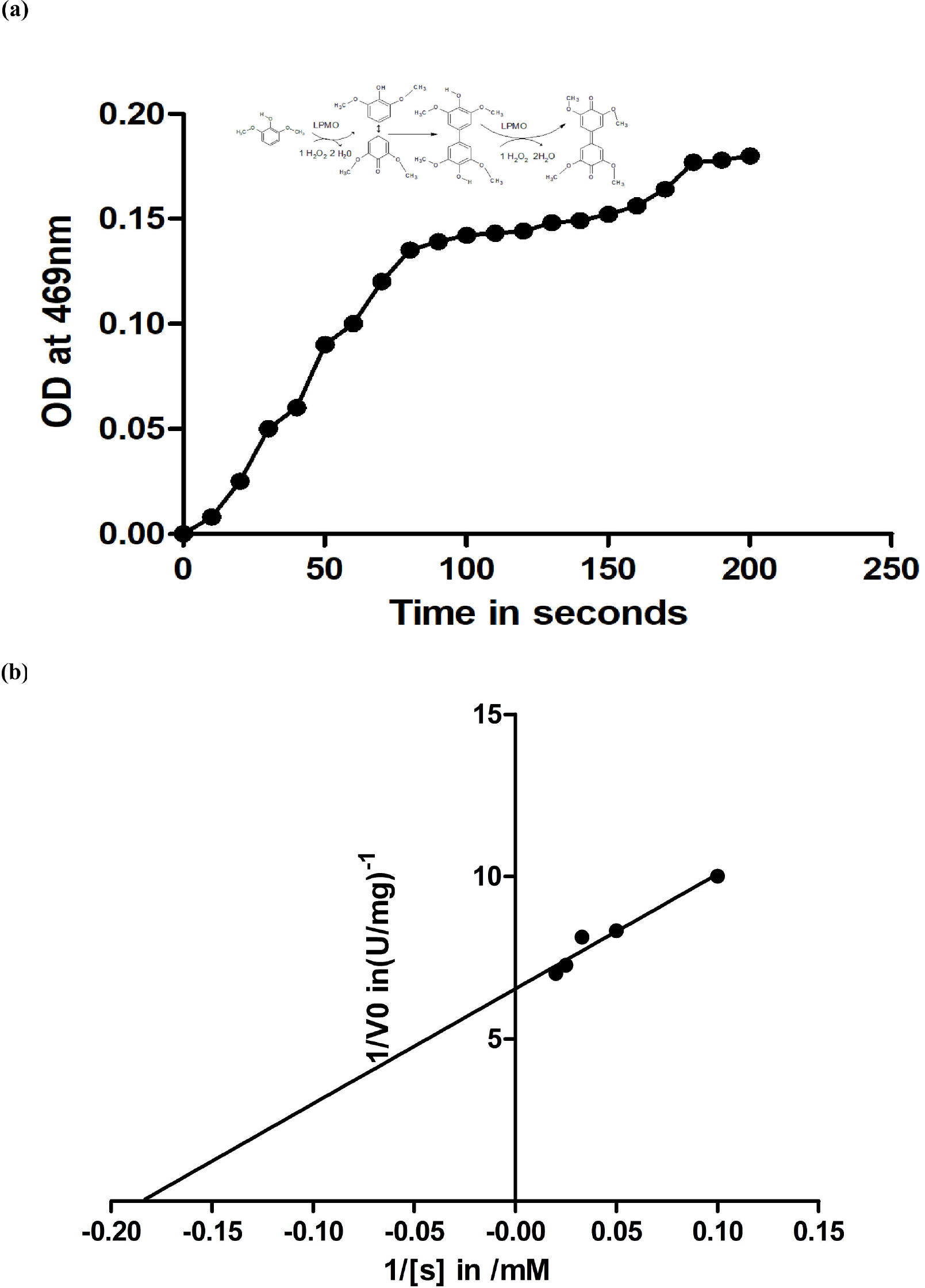
Enzyme kinetics studies of *Af*LPMO16 with 2,6-DMP (mean values are plotted). **(a)** Chemical reaction to convert 2,6DMP to 1-coerulignone; OD at 469 nm vs. time plot. **(b)** LB plot or 1/v vs 1/[s] plot.

**Table 1:** Enzyme kinetics of *Af*AA16 with 2,6, DMP as a substrate.

| Enzyme Kinetics Parameter | Values |
| --- | --- |
| $V_{\max}$ in U/mg | 0.153 |
| $K_m$ in mM | 5.4 |
| $K_{cat}$ in $\text{min}^{-1}$ | 277.67 |

### In-silico analysis for substrate specificity

The *Af*LPMO16 contains 19 amino acids long N-terminal signal peptide before His1 catalytic domain (1-169aa), and C terminal Serine rich region (170-271aa) (Fig. 3a). This N-terminal sequence is one of the marker features of fungal LPMOs, but this serine-rich C-terminal or linker is a feature of AA16 family. It also lacks the CBM1 module or glycosylphosphatidylinositol (GPI) anchor, like other AA16 LPMOs [19]. *Af*LPMO16 also has conserved Histidine at 1^st^ and 109^th^ positions, which are mainly involved in copper binding, the signature characteristic of LPMOs. There are other conserved sequences like Gly, Pro, Asn, Cys, Try, Tyr, Leu, and Asp, including GNV(I)QGELQ motif (Fig. 3b) The fully conserved sequences (highlighted with red background) are the marker amino acids represent the LPMOs. The partially conserved sequences (within the blue boxes) are the marker of different auxiliary families (Fig. 3b). The sequence alignment studies of AA16 family (including *Af*LPMO16) with other families (AA9, AA10, and AA11) of LPMOs suggested (Fig. S1) a co-relation between AA10 family and AA16 LPMOs. The substrate-binding motif in the L2 loop of cellulose active LPMO10 has some similarities with AA16 L2 loop motif (marked with black box) and cellulose active motif (Fig. 3b). In AA16 LPMOs the conserved motif in L2 loop GNI(V)QGEL the region is replaced by YNWFG(A)NL for C1 oxidizing AA10 LPMOs, which are also cellulose active. The previous study suggests that the amino acids (Y79, N80, F82, Y111, and W141) in loop L2 take part in substrate specificity for LPMO 10, and mutations (Y79, N80D, F82A, Y111F, W141Q) alter the specificity of the substrate from chitin to cellulose [37]. In *Af*LPMO16, the corresponding amino acids GNQYR (Fig. 3b) (marked with black arrows), some amino acids from these positions (N & Y) are also present in cellulose-active AA10 LPMOs. Hopefully, the polar amino acids (Q & R) are charged and may interact with chitin due to electrostatic interaction. Alternatively, there are high chances that few mutations in these amino acids may help *Af*LPMO16 to interact with chitin. Further, in chitin active LPMOs, more than 70% residues of the motif (Y(W)EPQSVE) are polar, including two negatively charged Glu (E). In cellulose active LPMOs, 70% residues of the motif (Y(W)NWFGVL) are hydrophobic [38]. In contrast, in *Af*LPMO16, 70% residues are polar, including one negatively charged Glu (E), one hydrophobic Tyr (Y), and others are neutral. The presence of polar residue and negative charged Glu (E) suggests that *Af*LPMO16 may bind to chitin. Electrostatics interaction between the substrate and enzyme active site plays a pivotal role in substrate binding. The electrostatic potential surface at the catalytic site of the *Af*LPMO16 was found unchanged or slightly positive-charged at pH 6.0 (Fig. 3c) (Marked in the figure). The electrostatic interaction study suggests that the *Af*LPMO16 may also bind to cellulose [52].

**Figure 3.**
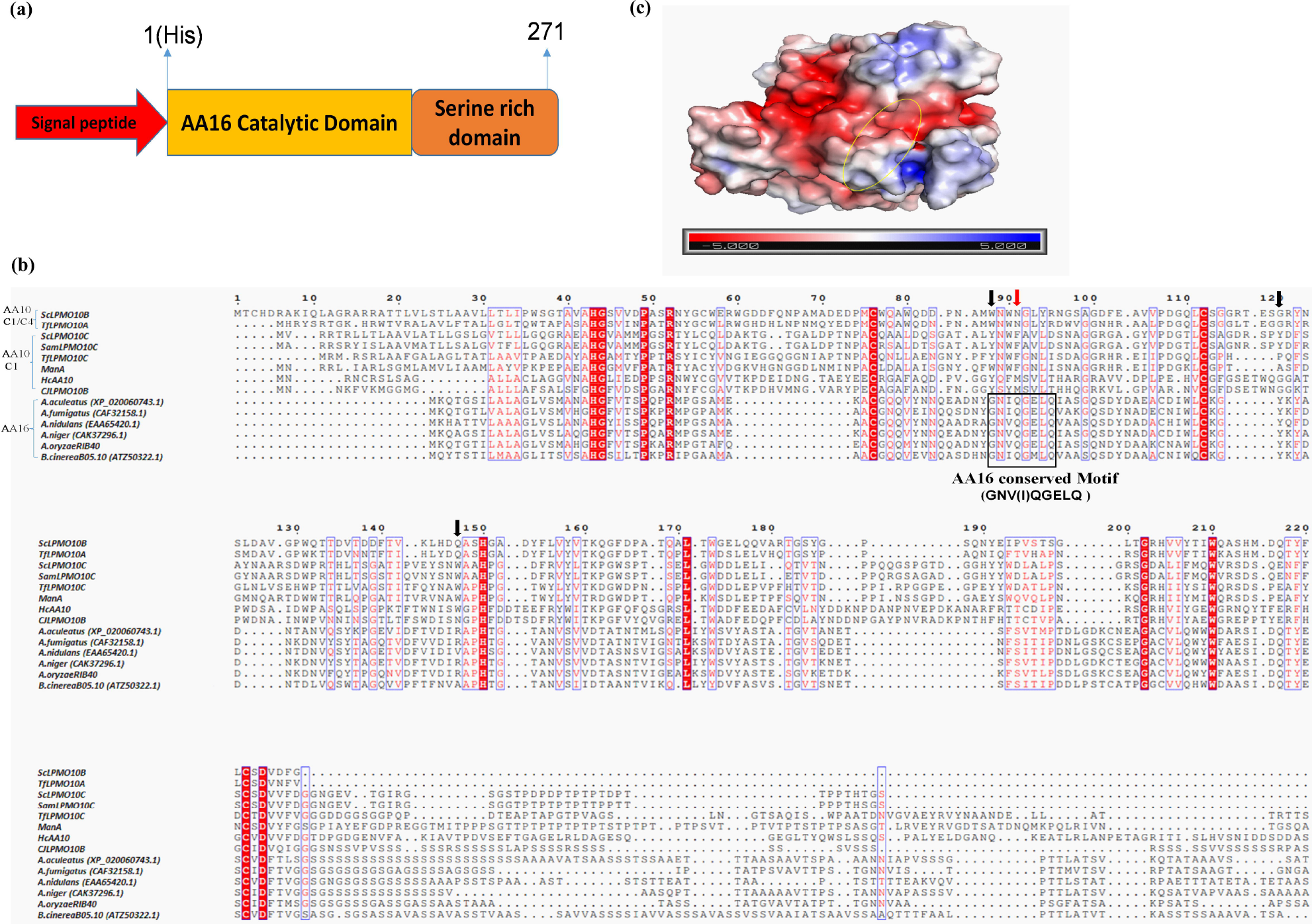
*In silico* analysis of *Af*LPMO16. **(a)** Schematic diagram of *Af*LPMO16; signal peptide: 19 amino acids, catalytic domain: 1-169 amino acids, and a serine-rich domain: 169-271 amino acids. **(b)** Multiple sequence alignment of AA16 LPMOs, C1 oxidizing, and C1/C4 oxidizing AA10 LPMOs: Conserved sequences are highlighted. The red arrow indicates the amino acid responsible for regioselectivity; the Black arrow represents the amino acid responsible for substrate specificity, the black box represents the AA16 conserved motif. **(c)** The electrostatic surface potential of *Af*LPMO16 model structure at pH6.0, blue and red color represents positive and negative potential surface respectively. The area surrounded by the ring represents the catalytic site.

### Regioselectivity of AfLPMO16

Amino acids on the substrate-binding surface determine the oxidative regioselectivity of LPMOs [29]. Sequence comparison and mutation studies revealed that the conserved amino acids near the catalytic center in C1 and C1/C4 oxidizing AA10 and AA9 LPMOs are responsible for regioselectivity. In the case of C1/C4 oxidizing AA10, the amino acid Asn85 near the catalytic center is responsible for C4 oxidizing activity. Alteration of this amino acid (N85F) diminished the C4 activity and produced only C1 oxidized product [39]. In C1 oxidizing AA9 LPMOs, hydrophobic amino acids Phe and Tyr are conserved in addition to Asn. While in C1 oxidizing AA10 LPMOs, the Phe amino acid has replaced the corresponding Asn site (Fig. 3b)(marked with red arrow). The Phe is also parallel to the substrate-binding surface [47]. In AA16, the corresponding Gln (Q) may be parallel to the substrate-binding region (Fig. 3b). The function of conserved Gln (Q) is not clear. However, this polar amino acid has a similar side chain with polar Asn (N). The axial distance between the conserved amino acid and copper catalytic center is another crucial factor for regioselectivity. The C1/C4 oxidizing AA10 LPMOs have more open or wider axial gaps than C1 oxidizing AA10 LPMOs [39]. Here the distance between Gln56 and His20 is 7.7Å, and the distance between Gln56 and Cu catalytic center is 11.1Å. In the absence of the AA16 structure (crystal or model), we cannot compare the lengths; nevertheless, this distance may play a key role in regioselectivity.

### Phylogenetic tree construction and analysis

The sequential and functional relationship of AA10 and AA16 LPMOs has been discussed, but phylogenetic studies based on the sequence similarity give an evolutionary origin. Based on sequence comparison, *Af*LPMO16 is evolutionarily closer to the LPMO of *Aspergillus fisheri* (91% sequence homology). The constructed phylogenetic tree contains two main clades and two subclades (Fig. 4). The first clade contains all AA10 LPMOs from bacterial species such as *Bacillus thuringiensis, Bacillus amyloliquefaciens, Streptomyces lividans*, and *Enterococcus faecalis*. The second clade includes all fungal AA10 and AA16LPMOs, mainly belongs to *Aspergillus*, and *Penicillium* species in which AA16 LPMOs are mostly from *A*.*niger, A*.*fumigatus, A*.*fisheri, Aspergillus kawachii* (Fig. 4).

**Figure 4.**
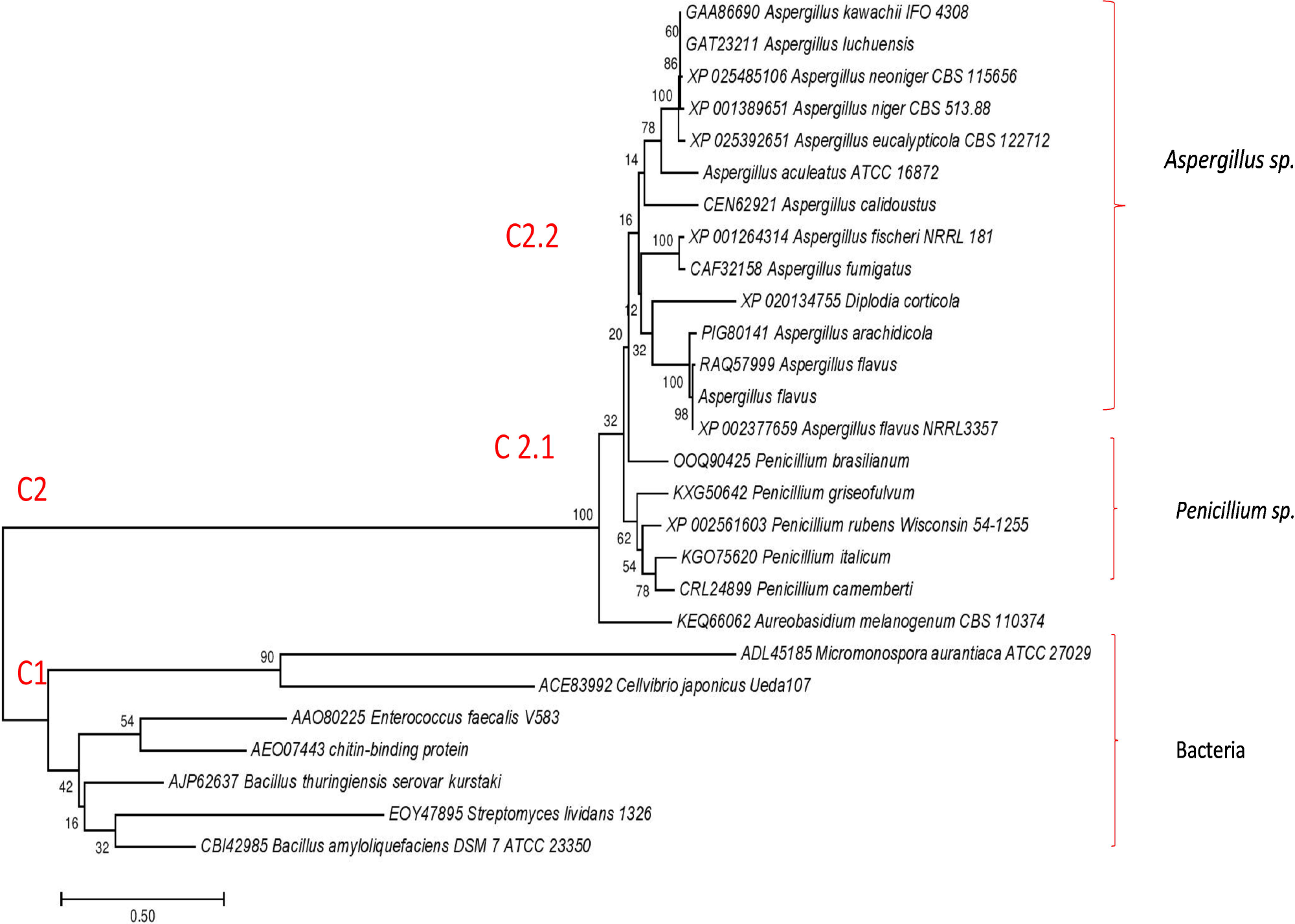
Phylogenetic relationship of *Af*LPMO16 with AA10 LPMOs. A neighbor-joining tree from MEGA showing C1(Bacterial) & C2(Fungal) clades and C2 clade further divided into C2.1 (*Penicillium* & other) & C2.2 (*Aspergillus*) subclades.

### Model structure prediction and molecular docking analysis

I-TASSER was used to predict the three-dimensional structure of the *Af*LPMO16. Most of the LPMOs have immunoglobulin-like distorted β-sandwich fold like structures, in which loops connect seven antiparallel β-strands with a different number of α-helix insertions (Fig. 5a). The final model has a β-sandwich structure connected by loops with two α-helices. The superimposition of the *Af*LPMO16 with other LPMO families like AA9, AA10, AA11, and AA13 showed that they share common antiparallel β-strands and helices with more loops, which indicate higher flexibility. Moreover, *Af*LPMO16 showed 1.2Å RMSD with AA11 (PDB Id: 4MAH) LPMO lower than the other LPMOs. So the 3D structure of *Af*LPMO16 suggests that it has more structural resemblance with AA11 LPMO. We also found one disulfide bond in *Af*LPMO16 between the Cys78-Cys186 amino acids, signature of thermo-stability (Fig. S2). The histidine brace amino acids, such as His20 and His109, participate in coordination bond with Cu ions. The surface of *Af*LPMO16 has an active site (Fig. 5b). The interaction studies with cellohexose suggest amino acids like Gln48, Gln181, Ser178, His109, His20, Asn54, Asp50, Tyr52, and Glu58 (Active enzyme starts with His1; so His20 will His1 and corresponding amino acids can be numbered accordingly) are in the active site and are involved in the interaction with the substrate (Fig. 5c). Molecular docking suggests that *Af*LPMO16 has a cellulose-binding surface (Fig. 5b & 5c). This study also suggests that the binding energy between *Af*LPMO16 and cellulose is −7.0 kcal/mol, which is highest compared to chitin (−5.5kcal/mol) and other polysaccharides.

**Figure 5.**
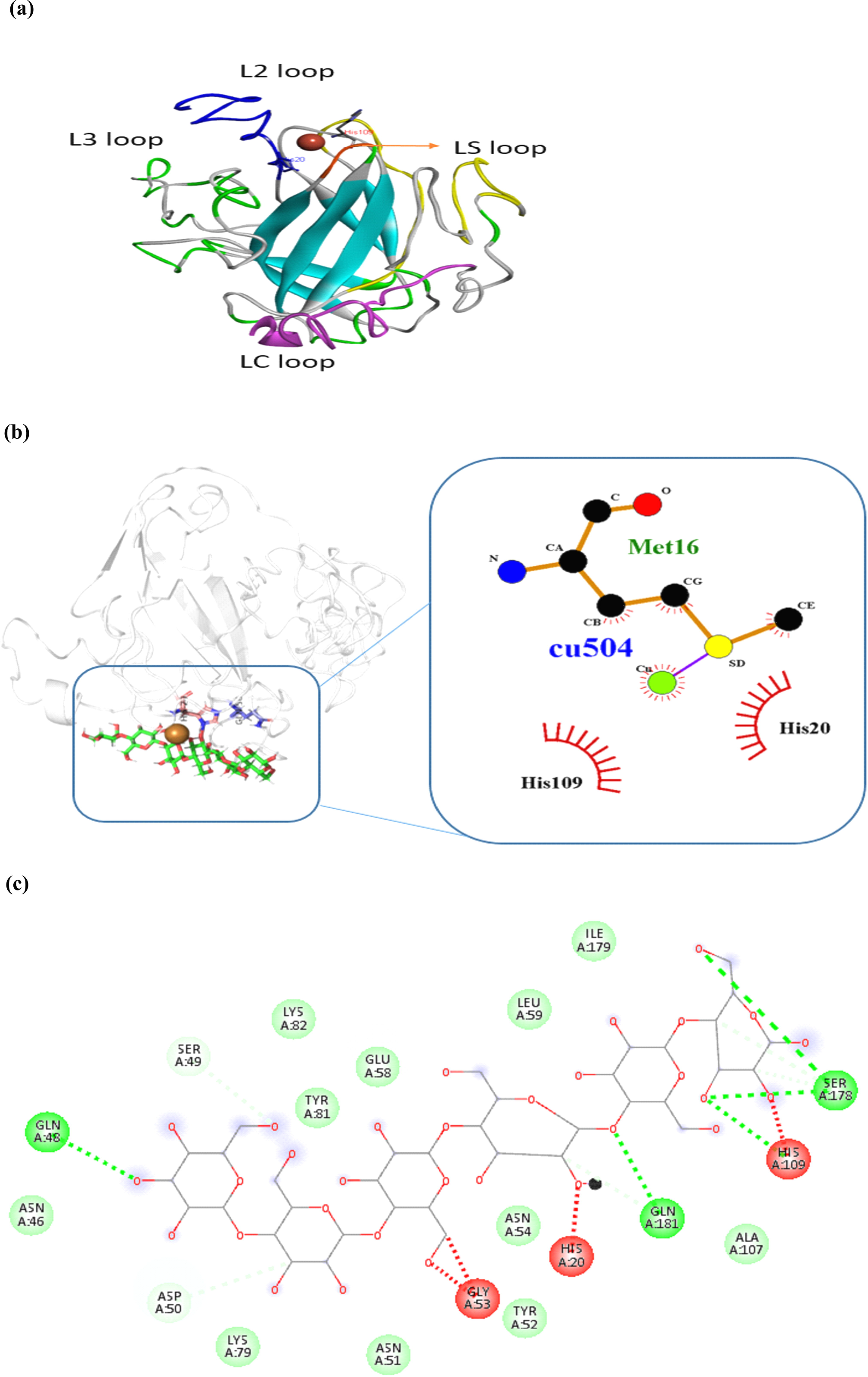
Model structure and molecular docking of *Af*LPMO16. **(a)** Predicted three-dimensional models of the *Af*LPMO16 showing functional loops LS(orange), L2(blue), L3(green), LC(magenta) loops surrounding the copper active site. **(b)** Histidine brace (His20, His109) of *Af*LPMO16 surrounding the copper metal. **(c)** Amino acids involved in substrate binding: Gln48, Gln181, Ser178, His109, His20, Asn54, Asp50, Tyr52, Glu58

### Polysaccharides depolymerization by AfLPMO16

*Af*LPMO16 showed efficient depolymerization activity on both CMC and PASC (Fig. 6a & 6b). We quantified the amount of reducing sugar released by enzymatic degradation. When incubated CMC with increasing concentrations of the enzyme, the amount of product (reducing sugar) increased with the increase of *Af*LPMO16 concentration (Fig. 6a). When we added 50µg of the enzyme, nearly 0.05mg/ml of reducing sugar was released. For 100µg of the enzyme, the product was nearly 0.136mg/ml, and for 200µg of the enzyme, the amount of product released was approximately 0.356mg/ml (Fig. 6a). This result indicates the polysaccharide (CMC) depolymerization activity of *Af*LPMO16.

**Figure 6.**
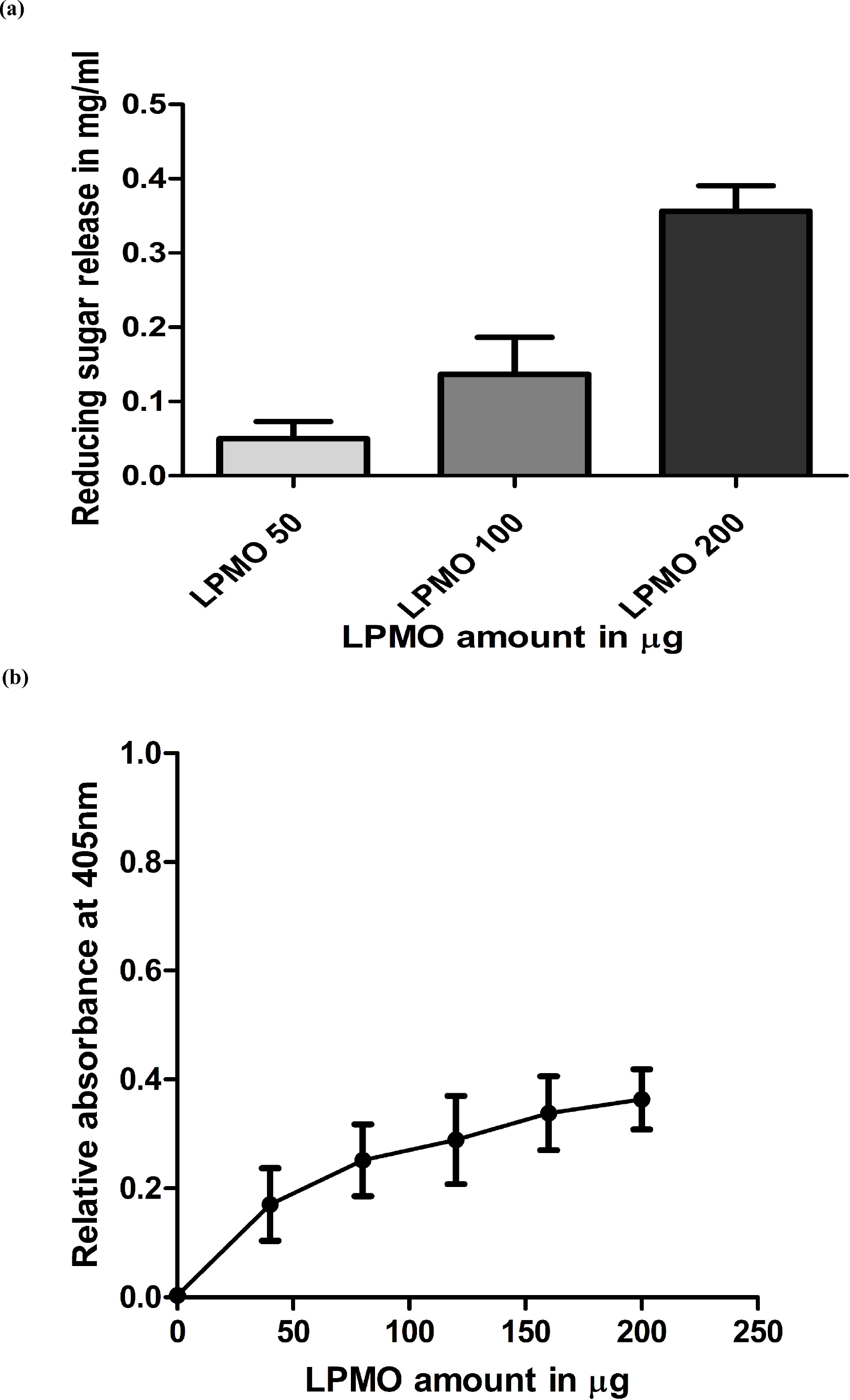
Polysaccharides degradation activity of *Af*LPMO16. **(a)** CMC depolymerization: estimation of reducing sugar with the increasing amount of *Af*LPMO16. **(b)** PASC hydrolysis: relative absorbance at 405nm vs. *Af*LPMO16 quantity plot. Results are the mean value of the minimum three experiments. The bar represents the standard deviation (SD)

Further, we used insoluble PASC as a substrate and incubated with an increasing concentration of *Af*LPMO16, and determined the relative absorbance of PASC with the growing amount of enzyme. The enzyme degrades the polysaccharide (substrate) into smaller polysaccharide units (monosaccharides, disaccharides, etc.), which are soluble and make the reaction mixture clearer. Therefore, it leads to a decrease in the absorbance resulting increment in relative absorbance [40]. Ultimately we will find a graph where relative absorbance increase with increasing concentration of *Af*LPMO16. Hence In this experiment, we found a rise in relative absorbance concerning the untreated substrate with a high concentration of enzyme *Af*LPMO16 (Fig. 6b). The graph (Fig. 6b) showed that 0.17 absorbance difference concerning untreated substrate when we used 50µl (concentration 0.8µg/µl) of the enzyme. The difference in absorbance steadily increased with the escalation of enzyme concentration (200µl of the enzyme at the concentration of 0.8µg/µl the relative absorbance reached nearly 0.36). Hence these experiments confirmed the intrinsic polysaccharide degradation property of the *Af*LPMO16 like other LPMOs. In these experiments, we used the heat-inactivated *Af*LPMO16 and ascorbic acid-deficient set to verify these results (data not shown).

### Pre-treated lignocellulosic biomass and cellulose hydrolysis with simultaneous treatment of AfLPMO16 and commercial cellulase

There are two modes of action to show the synergy or boosting effect of LPMO while using with cellulase-sequential assay and simultaneous assay. In the sequential assay, LPMO should add a prior time limit to cellulase. And in the simultaneous assay, both the enzymes LPMO and cellulase are being used together to the substrate. In this study, we chose to perform a simultaneous assay for two reasons; simultaneous assay shows better synergy or boosting in crystalline cellulose [41] than sequential one. Furthermore, we aimed to check the synergy or stimulating activity of commercial cellulase by *Af*LPMO16 so that it may include in the cocktail for better depolymerizing action. Here the boosting effect of *Af*LPMO16 was studied with a commercial cellulase cocktail on both cellulose (Avicel) and lignocellulosic biomass (alkaline pre-treated rice straw). The alkaline pre-treatment has a beneficiary over acid pre-treatment in terms of hydrolysis yield [48]. The reason is that alkaline pre-treatment sufficiently removes the lignin [42], but it preserves hemicelluloses [43]. When incubating Avicel with *Af*LPMO16 and cellulase, the amount of reducing sugar released was almost double compared to Avicel incubated with either cellulase alone or cellulase along with heat-inactivated *Af*LPMO16 (Fig. 7b). A similar kind of boosting effect we observed in every time point from 5 hrs to 72 hrs. We also found the synergistic impact of *Af*LPMO16 in lignocellulosic biomass transformation to fermentable sugar (Fig. 7a). When we incubated the alkaline pre-treated rice straw with 100 µg and 200µg of *Af*LPMO16 along with cellulase, almost 1.7 fold and slightly above 2-fold of reducing sugar were released respectively compared to lignocellulose incubated with either cellulase alone or cellulase along with heat-inactivated *Af*LPMO16 (Fig. 7a) suggests the enhancement is dependent on the amount of auxiliary enzyme *Af*LPMO16. For further elaboration of the synergistic effect of *Af*LPMO16, another set of reactions prepared where the biomass was treated with an increasing concentration of only *Af*LPMO16. A minimal amount of hydrolysis activity was there, nearly 0.04 mg/ml to 0.06 mg/ml, reducing sugar quantified for *Af*LPMO16 treated biomass (Fig. 7c). This hydrolysis activity of *Af*LPMO16 alone is negligible compare to only cellulase treated biomass.

**Figure 7.**
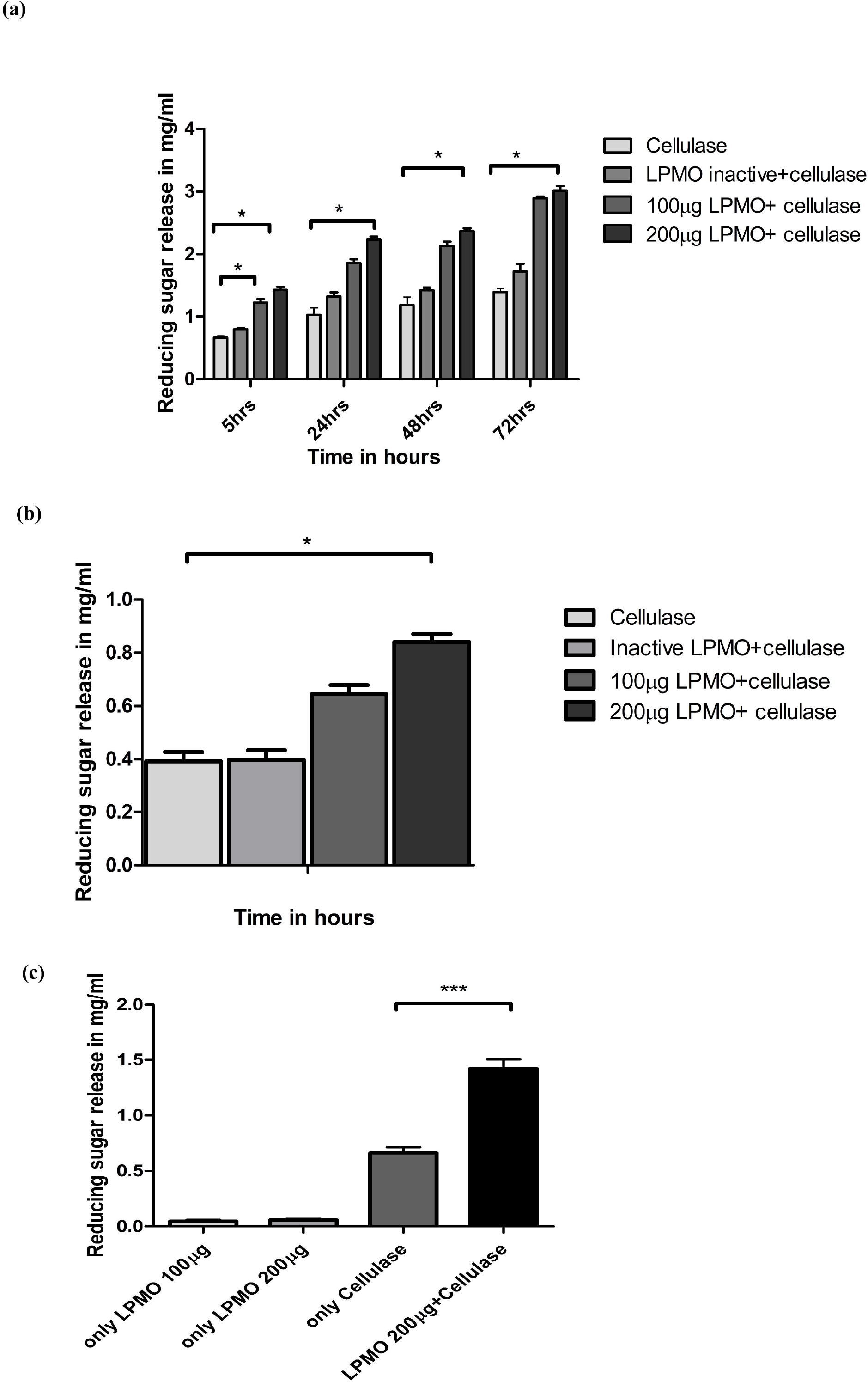
Boosting effect of *Af*LPMO16. **(a)** Hydrolysis of alkali pre-treated rice straw: light-grey bar indicates only cellulase and deep-grey indicates heat inactive *Af*LPMO16 with cellulase, dark-grey and black bar indicates cellulase along with two different quantity of *Af*LPMO16. **(b)** Avicel hydrolysis: reducing sugar estimation. Light-grey bar indicates only cellulase and deep-grey indicates heat inactive *Af*LPMO16 with cellulase, dark-grey and black bar indicates cellulase along with two different quantities of *Af*LPMO16. **(c)** Synergistic effect: light-grey bars indicate biomass hydrolysis by two different concentrations of *Af*LPMO16; dark-grey bar indicating the only cellulase treated biomass and black bar indicating combined treated biomass with *Af*LPMO16 & cellulase. Error bars represent the standard deviation of experiments ran in triplicate. The different number of asterisks (*) indicate a significant difference between glucose release in the presence of *Af*LPMO16 by one-way ANOVA followed by Student’s t-test (P<<0.05).

Nevertheless, the simultaneous use of *Af*LPMO16 and cellulase enhances the hydrolysis activity two-fold compared to the only cellulase treated biomass (Fig. 7c). This result strongly indicates the synergistic effect of *Af*LPMO16 with cellulase. All these results confirmed the boosting effect or synergistic effect of *Af*LPMO16 on the hydrolytic activity of cellulase for both cellulosic and lignocellulosic biomass degradation. So far highest synergistic effect was reported by AA9 (Table 2), which is less than two-fold [44, 45].

**Table 2:**
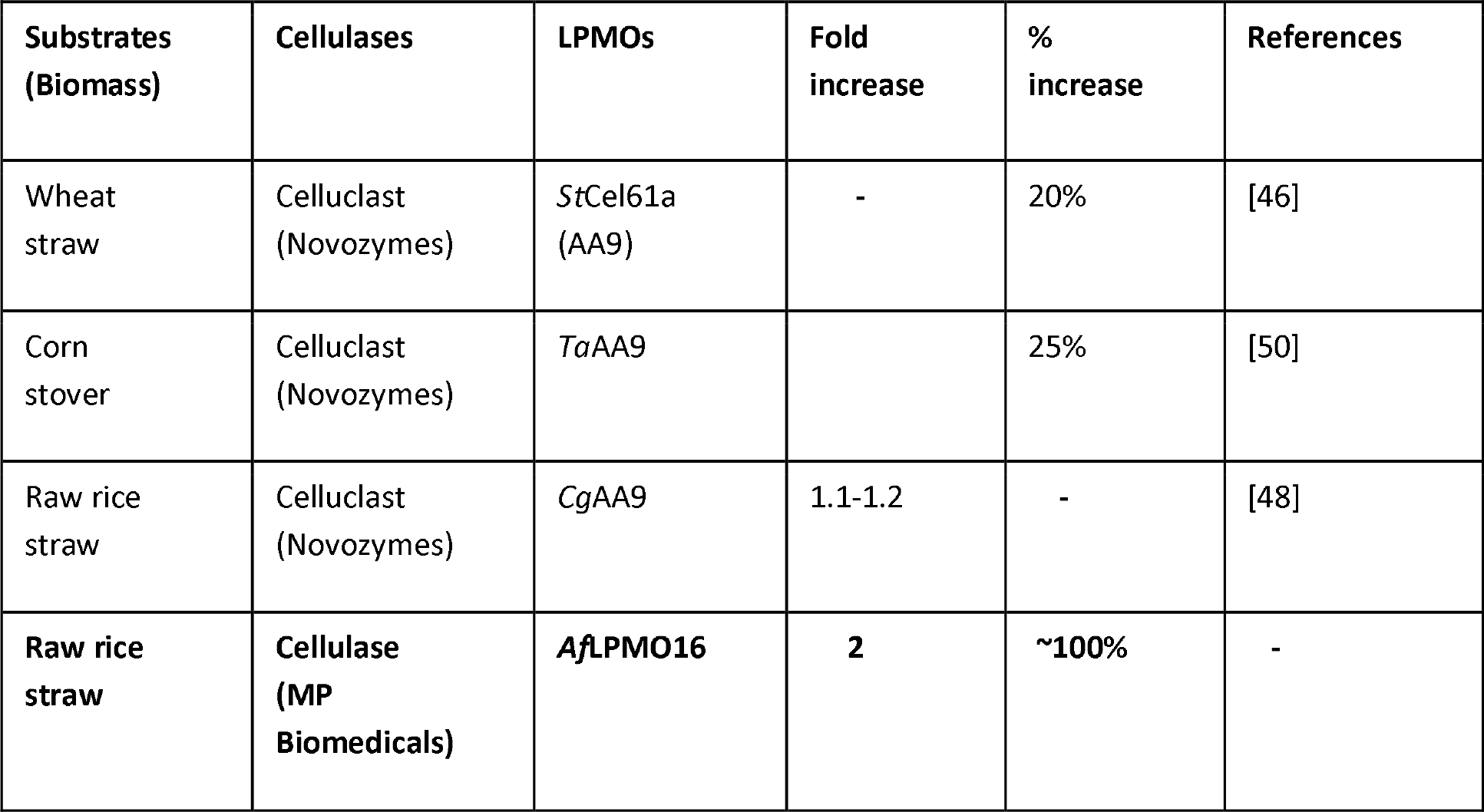
Lignocellulosic biomass hydrolysis enhancement by LPMOs

## Discussion

The gene was cloned in pPICZαA vector under the control of *AOX1* promoter by following the same strategy developed for *Aa*AA16 and PMO9A_MLACI [19, 26]. The nucleotide sequence of *Af*LPMO16 was codon-optimized for *Pichia pastoris*. The recombinant protein containing a C-terminal polyhistidine tag was produced in flasks in the presence of trace metals, including copper, and purified from the culture supernatant by immobilized metal ion affinity chromatography (IMAC: Ni-NTA affinity chromatography), following the same protocol used for *Aa*AA16 [19]. We were successful in producing the active *Af*LPMO16 in *P*.*pastoris* X33 (Fig. 1) in a shake flask. Despite the chance of N-terminal modification in shake flask culture instead of bioreactor culture [19], the amount of active enzyme obtained in shake flask was sufficient for characterization. The enzyme activity determined by 2,6-dimethoxyphenol concerning the heat-inactivated enzyme and without ascorbic acid as negative controls (data not shown). The enzyme activity suggests the successful production of active protein (Fig. 2a), and interestingly, the initial reaction rate is faster compared to later time span. Lytic polysaccharide monooxygenase (LPMO) releases a spectrum of cleavage products from their polymeric substrates cellulose, hemicellulose, or chitin. The correct identification and quantitation of these released products is the basis of MS/HPLC-based detection methods for LPMO activity, which is time taking and is required specialized laboratories to measure LPMO activity in day-to-day work. A spectrophotometric assay based on the 2,6-dimethoxyphenol can accurately measure the enzymatic action and can be used for enzyme screening, production, and purification, and can also be applied to study enzyme Kinetics [21]. Thus it is swift, robust for biochemical characterization, and also accurately determines the active enzyme.

Sequence analyses indicating that the *Af*LPMO16 has some signature characteristics for both cellulose and chitin-binding and both C1 and C1/C4 oxidizing activity. However, experimental confirmation is required to establish the presence or absence of any chitin-binding nature and C1/C4 oxidizing capability of *Af*LPMO16. The constructed phylogenetic tree (Fig. 4) suggests that the fungal AA10 and AA16 LPMOs are more likely to come from a common ancestor. Molecular docking study suggests that *Af*LPMO16 has the highest affinity towards cellulose among the known substrates, based on the binding energy. The binding energy between cellulose and *Af*LPMO16 is −7.0 Kcal/mol, which makes thermodynamically strong binding between enzyme and substrate (Fig. 5b & 5c) compared to other substrates. The LPMOs are essential for their auxiliary activity and polysaccharide degrading property. We observed polysaccharide depolymerizing activity on carboxymethyl cellulose (CMC) and phosphoric acid swollen cellulose (PASC) (Fig. 6a & 6b). Due to its auxiliary activity, it enhances the action of the cellulase enzyme for the degradation of cellulose and lignocelluloses [49]. The only identified AA16 family, the *Aa*AA16, showed a sequential boosting effect with *T. reesei* CBHI on nano-fibrillated cellulose (NFC) and PASC. The *Aa*AA16, the recent addition of the AA16 family of LPMO in the CAZY database, showed synergism with the CBH1 for the degradation of cellulose [19]. However, *Aa*AA16 study did not deal with the biomass hydrolysis boosting effect of the AA16 family. The boosting result is most important in the technical aspect for enhancing the activity of the cellulase cocktail. LPMO enzyme has earned much research interest due to their synergistic effect or boosting effect on cellulase enzyme [45]. *Af*LPMO16 showed a boosting impact on cellulose and lignocellulose hydrolysis (Fig. 7a & 7b). The synergism of *Af*LPMO16 has shown in (Fig. 7c), where the only *Af*LPMO16 and only cellulase treated biomass hydrolysis activity is low compare to the combined effect of these two enzymes. The simultaneous use of *Af*LPMO16 and cellulase enhances nearly two-fold biomass hydrolysis compare to the only cellulase treated biomass hydrolysis. This enhancement of two-fold biomass hydrolysis is higher than that of other LPMO families [50]. However, the synergy or boosting effect depends on many factors such as pre-treatment [51], the lignin content of lignocelluloses and acting cellulase [46]. Still, over 50% enhancement suggests intense demands on inclusion on cellulase cocktail. However, the mechanism of synergism with the cellulase enzyme complex is poorly understood. The probable explanation of such a boosting effect could be that the cellulosic biomass is partially depolymerized by the LPMO, which gives further access to the cellulase enzymes.

## Conclusion

In concluding remark, *Af*LPMO16 is the second report of the AA16 family of LPMO, but for the first time, we have characterized the AA16 family biochemically and structurally. *In-silico* sequence analysis, structure analysis, and molecular docking studies suggest some unique characteristics of the *Af*LPMO16, like cellulose-binding ability, chances of chitin-binding, and C1 and C4 oxidizing property. Further studies, including the engineering approach, are required to confirm these characteristics. Nevertheless, the most crucial aspect of *Af*LPMO16 is the significant boosting effect on commercial cellulase cocktail in lignocellulosic biomass conversion, and that suggests its importance in the bioethanol industry.

## Materials and Methods

### Sequence analysis and Phylogenetic analysis

*Af*LPMO16 sequence (CAF32158.1) was obtained from NCBI, and the sequence was further confirmed from the *Aspergillus* genome database (http://www.aspgd.org/). To avoid interference from the presence or the absence of additional residues or domains, the signal peptides, and C-terminal extensions were removed before the alignment. Homology sequence alignment was performed by the BLAST [22]. Clustal Omega [23] was used for multiple sequence alignment. The sequence alignment was edited with Espript for better visualization. Pymol [24] and MEGA7 [25] were used to construct a phylogenetic tree after sequence alignment. To build the phylogenetic tree, the sequences of twenty-seven (27) LPMO genes (edited to remove N-terminal signal sequence, C-terminal extension or GPI anchor, CBM1 module) were taken from different species belong to AA10 and AA16 family of LPMOs. The neighbor-joining tree was constructed with 1000 bootstrap replications.

### Cloning of AfLPMO16

*Aspergillus fumigatus* NITDGPKA3 was grown on CMC agar media containing 2% CMC, 0.2% peptone, 2% agar in basal medium (0.2% NaNO_3_, 0.05%KCl, 0.05%MgSO_4_, 0.001%FeSO_4_, 0.1%K_2_HPO_4_). The fungal biomass was then milled in a pestle and mortar followed by rapid overtaxing in solution with an appropriate lysis buffer for proper lysis of the cell. Genomic DNA was isolated from the fungal biomass using the DNA extraction buffer (400mM Tris-HCl, 150mM NaCl, 0.5M EDTA, 1%SDS) and followed by Phenol, chloroform and isoamyl alcohol (25:24:1) extraction. The final pellet was washed with 70% alcohol, air-dried, and dissolved in sterile water. *Af*LPMO16 gene was amplified by polymerase chain reaction (PCR). The codon-optimized gene for *Pichia pastoris* was inserted into the pPICZαA vector (Invitrogen Carlsbad, California, USA). The gene was cloned with the native signal sequence and 6x His-tag at the C-terminal [26]. The cloning was done by following the same protocol as *Aa*AA16 and PMO9A_MLACI [19, 26]. The vector (pPICZαA) containing the *Af*LPMO16 gene was linearized by Pme1 (New England BioLabs) and transformed to *Pichia pastoris* X33 competent cells. The Zeocin resistant transformants were picked and screened for protein production. The cloned gene was further confirmed by sequencing and the sequence submitted to GenBank (GenBank accession No. MT462230).

### Expression and purification of AfLPMO16

The positive colonies were selected on YPDS (Zeocin: 100μg/ml) plates. The positive transformants were further screened by the colony PCR and expression studies. Protein expression was carried out initially in BMGY media containing 1ml/L *Pichia* trace minerals 4 (PTM4) salt (2g/L CuSO_4_·5H_2_O, 3g/L MnSO4·H_2_ O, 0.2g/L Na_2_MoO_4_·2H_2_O, 0.02g/L H_3_BO_3_, 0.5g/L CaSO_4_·2H_2_O, 0.5g/L CoCl_2_, 12.5g/L ZnSO_4_·7H_2_O, 22g/L FeSO_4_·7H_2_O, NaI 0.08g/L, H_2_SO_4_ 1mL/L) and 0.1 g/L of biotin. Then after 16 hours, *Pichia* cells were transferred into BMMY medium (PTM4 salt) with continuous induction by the addition of 1% methanol (optimized) every day (after every 24 hours) for three days. After three days, the culture media was spun down (8,000rpm for 10mins) at 4^0^C. The pellet was discarded, and the media was collected. The protein was precipitated from the media by ammonium sulfate precipitation (90% saturation). The pellet was redissolved in Tris buffer (Tris-HCl 50mM pH-7.8, NaCl-400mM, Imidazole-10mM). The recombinant protein was purified by immobilized ion affinity chromatography (Ni-NTA affinity chromatography)[27], followed by dialysis with 50mM phosphate buffer, pH 6.0. We followed the expression and purification procedure, same as *Aa*AA16 [19]. The yield of the purified protein was almost 0.8 mg/ml. The concentration was measured by Bradford assay, and BSA was used for standard concentration. The protein was separated by SDS-PAGE using 12% acrylamide in resolving gel(dH_2_O-3.6 ml, Acrylamide+Bisacrylamide – 4.0 ml, 1.5M Tris-2.6 ml, 10%SDS-0.1 ml, 10% APS-0.1 ml, TEMED-0.01 ml; for 10 ml), stained with coomassie blue, and the purified protein band was also confirmed by Western blot analysis by using an anti-His antibody (Abcam).

### Biochemical assays of AfLPMO16

#### Biochemical characterization of AfLPMO16

2,6 DMP (2,6-dimethoxyphenol) was used as a substrate for *Af*LPMO16 in this study. The reaction was done in phosphate buffer (100mM pH 6.0) containing 10mM 2,6-dimethoxyphenol, 5μM hydrogen peroxide, and 50μg of purified *Af*LPMO16 at 30□C. The amount of product 1-coerulignone was measured by spectrophotometer using the standard extinction coefficient (53200M^-1^cm^-1^) and Lambert-Beer law. For kinetic assay different 2,6-dimethoxyphenol concentrations (1mM, 5mM, 10mM, 20mM, 25mM, 30mM, 40mM, 50mM, 70mM and 100mM) were used. The kinetic parameters were calculated based on the Line-weaver-Burk plot (LB plot). One unit of enzyme activity is defined as the amount of enzyme which releases 1μM of 1-coerulignone (product) per minute in standard reaction condition.

#### Polysaccharides depolymerization by AfLPMO16

Different cellulosic compounds such as PASC, avicel®PH-101 (SIGMA), and carboxyl methylcellulose (CMC) was used. We used 1% Avicel®PH-101 (SIGMA) (crystalline cellulose) and 1% CMC (Carboxyl methylcellulose sodium salt) with different concentrations of purified *Af*LPMO16 for different incubation time. Reducing sugar was determined by Dinitro salicylic acid (DNS) assay. For PASC assay, we used 0.25% PASC and incubated with increasing concentration of *Af*LPMO16 for 6 hours and measured the OD after 6hrs of incubation and plot the relative absorbance ([OD of *Af*LPMO16 treated PASC]-[OD of untreated substrate]) with enzyme concentration [28].

#### Biomass and cellulose hydrolysis by cellulase and AfLPMO16

Cellulose and lignocellulose (alkaline pre-treated raw rice straw) [29] was used to determine the cellulose hydrolysis enhancing capacity. Rice straw was pre-treated with 5% NaOH (1:10 W/V ratio) at 120□C at 15Psi pressure for 1 hour, and sodium azide (20%) 10μl (per 10ml) was added at the reaction mixture to prevent any microbial contamination. The reaction was performed at 50□C, and the amount of reducing sugar was quantified after 5hours, 24hours, 48hours, and 72 hours by Dinitro salicylic acid (DNS) assay. 20μl of cellulase (commercial) (MP Biomedicals LLC) (5mg/ml) was used along with two different concentrations of *Af*LPMO16 125μl (100μg) and 250μl (200μg) [concentration 0.8mg/ml]. Reaction sets were prepared using the only cellulase, only *Af*LPMO16 with different concentrations, combined *Af*LPMO16 and cellulase and lastly, cellulase with inactivated *Af*LPMO16. *Af*LPMO16 was heat-inactivated by keeping at 100□C temperature for 30 minutes. Reducing sugar from each triplicate sets were quantified. In the case of cellulose degradation, 400μl (1%) of avicel (SIGMA) was incubated with 10μl of cellulase (commercial) (MP Biomedicals LLC) (5mg/ml). Reducing sugar was quantified after 5 hours of incubation. For these biochemical assays, we used 100mM phosphate buffer (pH-6.0), and heat-inactivated *Af*LPMO16 was taken as a negative control.

#### Molecular modeling and Molecular docking

I-TASSER [30] server was used to model the *Af*LPMO16. The final model was energy minimized by Gromacs software [31]. The Ramachandran plot [32] and Procheck [33] was used to evaluate the final model. For Metal Ion-Binding site prediction and docking server or MIB server (http://bioinfo.cmu.edu.tw/MIB/) were used to identify the copper (Cu) ion position. A molecular docking study was performed by the Autodock Vina [34] using MGL tools (Molecular graphics laboratory). The optimized substrate structures were prepared by Autodock vina and saved in PDBQT format. The grid size parameters used in this docking were 44, 46, 46, and grid center parameters used in this study were 49, 45, and 55. The genetic algorithm was also used for docking. Molecular interactions between enzyme and substrate were analyzed by the MGL tools [35]. The electrostatic potential surface of the *Af*LPMO16 is calculated by the APBS plugin available in Pymol at pH 6.0.

## Acknowledgments

MH is thankful to DBT, and SRD is grateful to DST Inspire for their fellowship. The authors are also thankful to DST-FIST grant of the Department of Biotechnology, NIT Durgapur.

## Authors’ contribution

MH and SM designed the research work. MH, BSK, and SM wrote the manuscript. MH performed biochemical assays. SRD performed *In-Silico* analysis. MH and KA analyzed the results. All authors read and approved the manuscript.

## Ethical approval

No human participants or animal is being used during the study.

